# Alzheimer’s Amyloid-Beta Intermediates Generated by Polymer-Nanodiscs

**DOI:** 10.1101/435396

**Authors:** Bikash R. Sahoo, Takuya Genjo, Michael Bekier, Sarah J. Cox, Andrea K. Stoddard, Magdalena Ivanova, Kazuma Yasuhara, Carol A. Fierke, Yanzhuang Wang, Ayyalusamy Ramamoorthy

## Abstract

Polymethacrylate-copolymer (PMA) encased lipid-nanodiscs (~10 nm) and macro-nanodiscs (>15 nm) are used to study Aβ_1-40_ aggregation. We demonstrate that PMA-nanodiscs form a ternary association with Aβ and regulate its aggregation kinetics by trapping intermediates. Results demonstrating reduced neurotoxicity of nanodisc-bound Aβ oligomers are also reported.

---

Among various proposed causes for the onset of Alzheimer’s disease (AD),^1,2^ the deposition of amyloid-β (Aβ) peptides, that are sequentially cleaved from an amyloid precursor protein in the brain, has remained as the fundamental hallmark for pathogenesis. The structural plasticity of unfolded Aβ monomers to adopt transient oligomers has been investigated to confer their neurotoxicity *in vitro*.^3,4^ The cell membrane has been shown to play crucial roles in modulating Aβ aggregation and redirecting aggregates to several intermediate states.^5,6,7^ Recent studies have further shown gangliosides (GM), sphingomyelin (SM) and cholesterol to be the major modulators of Aβ oligomerization.^8,9^ Although, recent studies proposed the catalytic activities of lipid-membrane on Aβ deposition and aggregation, the exact function of these lipids is not yet fully understood.^9,10^ Herein, we have investigated the roles of membrane composition in modulating Aβ’s aggregation using polymethacrylate (PMA) copolymer encased lipid-nanodiscs in an attempt to trap and characterize structure and toxicity of Aβ intermediates.

The formation of nanodiscs with different lipid composition was achieved by varying the lipid:PMA molar ratio (see methods in the supporting information). Dynamic light scattering (DLS) measurements depicted a hydrodynamic diameter ranging from ~ 8 to 10 nm for the targeted nanodiscs (Fig.S1a). For a comparative analysis and as per the suitability of PMA to form nanodiscs, we used 1,2-dimyristoyl-sn-glycero-3 lipids (broad phase transition temperature suitable for biophysical studies like NMR).^11^ The formation of mixed lipid-nanodiscs was further validated using ^31^P NMR experiments (Fig.S2a). We first observed the effects of PMA lipid-nanodiscs on Aβ_1-40_ aggregation kinetics by the thioflavin T (ThT) fluorescence assay (Fig.1). Results showed both lipid concentration and composition are important to regulate Aβ_1-40_ fibrillation. All nanodiscs showed no ThT binding in absence of Aβ_1-40_ in solution (Fig.S1b). Although the effect of nanodiscs on Aβ’s aggregation kinetics is complex (Fig.1), several trends can be observed. At 1:1 peptide:lipid molar concentration, all nanodiscs promoted aggregation irrespective of the lipid composition (Fig.1) with a significant decrease in the lag time (T_lag_).

**Figure 1.**
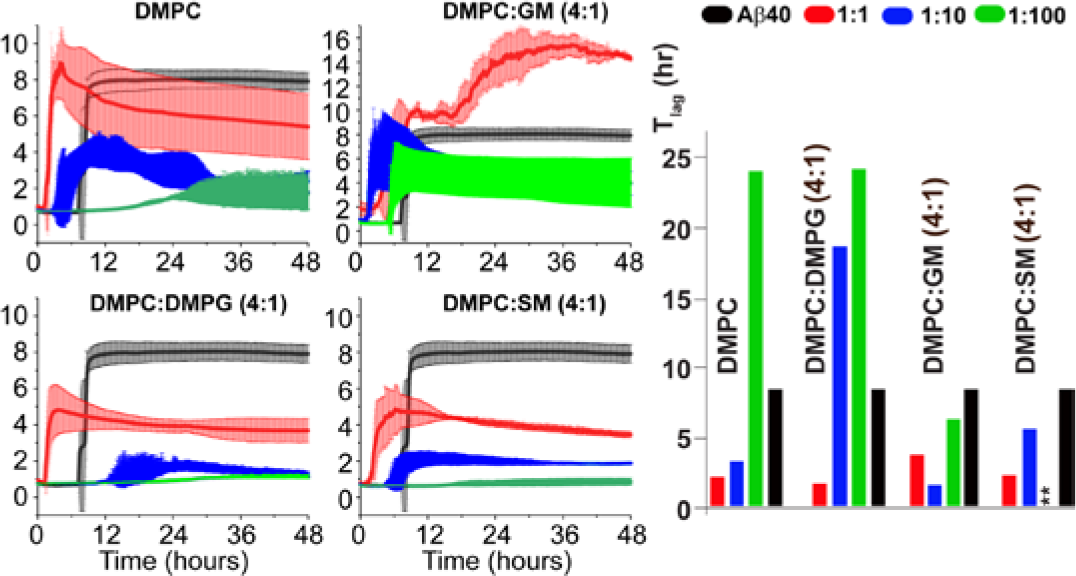
Aggregation kinetics of Aβ_1-40_ in presence of polymer lipid-nanodiscs. The aggregation kinetic profiles of Aβ_1-40_ (5 μM) was monitored by the changes in ThT fluorescence as a function of time to represent peptide alone (grey trace) and in the presence of varying concentrations (color traces for the indicated P:L ratios) of lipid-nanodiscs as shown on the top. GM, SM, DMPC and DMPG denote ganglioside, sphingomyelin, 1,2-dimyristoyl-sn-glycero-3-phosphocholine and 1,2-dimyristoyl-sn-glycero-3-phosphoryl-3'-rac-glycerol, respectively. The lag-time (T_lag_) is shown on the right. The asterisks indicate data not determined.

**Figure 2.**
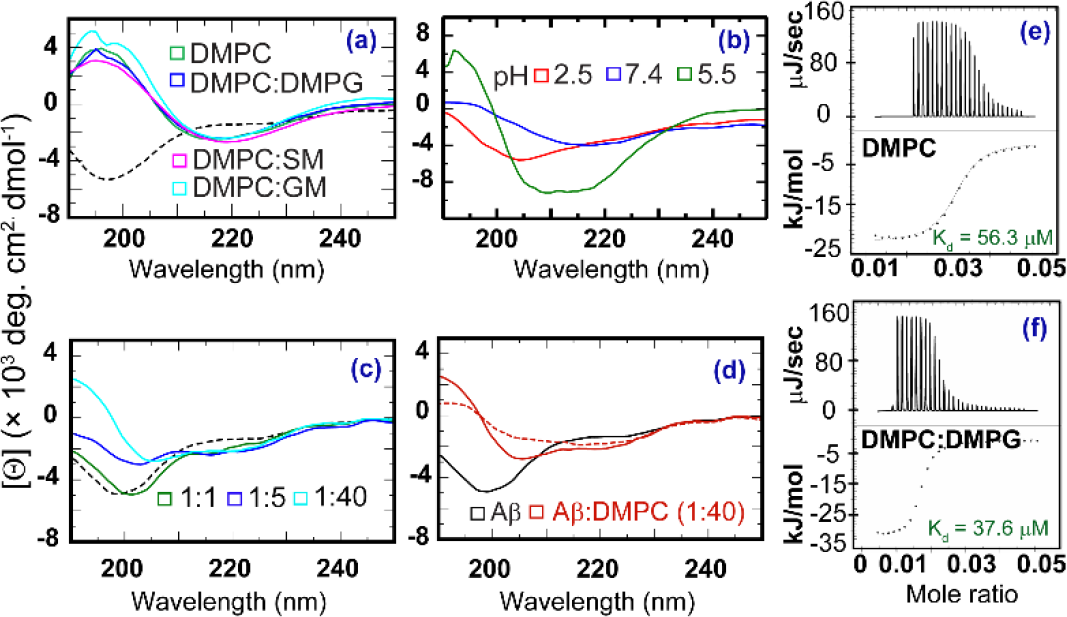
Conformational and kinetic analysis of Aβ_1-40_ binding to nanodiscs. (a) CD spectra of Aβ_1-40_ (25 μM) alone (dashed, black) and in the presence of different lipid-nanodiscs at peptide to lipid molar ratio of 1:10 (colored traces) measured at 25 °C. (b) pH dependent conformational changes in Aβ_1-40_ interacting with only PMA (red, blue) or PMA-encased DMPC nanodiscs (green) at a peptide to PMA molar ratio of 1:1. (c) CD spectra of Aβ_1-40_ (25 μM) interacting with DMPC macro-nanodiscs (diameter >16 nm) at the indicated peptide:lipid molar ratio (colored traces). (d) Secondary structure transition of Aβ_1-40_ due to binding to DMPC macro-nanodiscs as a function of time. The solid and dashed red lines denotes spectra measured at ~ 0 and 6 hours, respectively. (e-f) ITC thermograms showing the binding kinetics between Aβ_1-40_ and lipid-nanodiscs (DMPC or 4:1 DMPC/DMPG) at a peptide (20 μM) to lipid (400 μM) molar ratio of 1:20.

With increasing concentration of lipids (P:L=1:10), all nanodiscs except DMPC/DMPG (4:1) promoted Aβ_1-40_ aggregation (Fig.1, red traces). Remarkably, a substantial delay in Aβ_1-40_ aggregation was observed (except GM containing nanodiscs) by further increasing the lipid concentration (P:L=1:100). The differential activity between SM and GM1 nanodiscs on modulating Aβ_1-40_ aggregation kinetics are in agreement with a recent study that proposed counter protective roles of SM and GM1 on Aβ_1-40_ aggregation.^9^

Next, we monitored the effect of varying lipid composition on the conformation of Aβ_1-40_. Circular dichroism (CD) spectra of Aβ_1-40_ showed a rapid (< 5 minutes) structural transition from a random coil with spectrum minimum at ≈200 nm to a β-sheet conformation with spectrum minimum at ≈216 nm in all nanodiscs irrespective of the lipid composition (Fig.2a).

Interestingly, a previously reported Aβ_1-40_ oligomer (dimer, trimer or tetramer) also exhibited CD spectral with single minima between ≈210 nm and ≈216 nm^12,13^, which could indicate that the nanodiscs-bound β-sheet species could be an oligomer. However, unlike in the presence of nanodiscs, random-coil to β-sheet structural transition in solution occurs over several days. But, our ThT results show no fiber formation in Aβ_1-40_ within the CD measurement time i.e < 5 minutes for any of the lipid-nanodiscs (Fig.1). Therefore, the observation in the presence of nanodiscs suggests an induction of β-sheet-rich Aβ_1-40_ oligomers that are incompetent to bind ThT and unable to proceed the self-seeding reaction as seen in the presence of high lipid concentration. The fast aggregation observed at low P:L ratio may be attributed to the interaction of the peptide with the positively-charged PMA polymer-belt.

To explore the effect of PMA interactions with Aβ_1-40_ on the observed conformational transition, CD titration experiments were performed at varying pH. Conformational changes due to Aβ_1-40_ binding with PMA alone or with PMA-DMPC nanodiscs were measured at both acidic and basic conditions. We observed PMA at equimolar concentration of peptide in absence of lipids induced β-sheet conformation in Aβ_1-40_ at pH 7.4 where it carries a net charge of -2.9 (Fig.2b, blue trace). A decrease in the buffer pH from 7.4 to 2.5 reduces the electrostatic interaction between PMA and Aβ_1-40_ (net charge +6.7) resulting in a random-coil like Aβ_1-40_ conformation (Fig.2b, red trace). On the other hand, we observed a helical conformation of Aβ_1-40_ mixed with DMPC nanodiscs at pH=5.5 (the stability of nanodiscs were not affected at pH 5.5) where it carries nearly no charge (Fig.2b, green trace). The CD observation suggest that at physiological pH, the electrostatic interaction could be the driving force for the rapid structural conversion in Aβ_1-40_. Combining the CD and ThT results, we speculate that a concerted ternary association between the nanodisc (lipid and PMA-belt) and the peptidemodulates Aβ’s folding and aggregation and could be a useful tool to trap intermediate structures as we recently observed in peptide-based lipid-nanodiscs^14^.

The ternary association of PMA-nanodiscs with Aβ_1-40_ limits the interpretation of lipid directed Aβ_1-40_ aggregation. To diminish the effect of PMA and to understand the role of lipids alone in modulating Aβ_1-40_ aggregation at physiological pH 7.4, we designed macro-nanodiscs (diameter > 15 nm) using a lipid:PMA (w/w) ratio of 4:1 (Fig.S3a). CD spectra of Aβ_1-40_ in DMPC macro-nanodiscs (DMPC_md_) showed a lipid concentration dependent Aβ_1-40_ structural transition with a gradual change from random-coil to β-sheet conformation (Fig.2c). Aβ_1-40_ exhibited a mixed α-helix and β-sheet conformation for Aβ_1-40_:DMPC_md_ molar ratio of 1:5 or 1:40 and is in agreement with previous findings.^15^ At molar ratios (Aβ_1-40_:DMPC_md_) of 1:40, a time-dependent transition from α to β-sheet rich conformation was observed. CD spectral analysis using BeStSel^16^ predicted an α/β (%) structural content of 27.2/28.5 and 8.8/20.6 in Aβ_1-40_ in presence and absence of DMPC macro-nanodisc, respectively (Fig.2d). The random coil to α-helix change in the peptide structure further ensured the lipid dependent Aβ_1-40_ aggregation in macro-nanodiscs, unlike the observation from nanodiscs. The macro-nanodiscs comprising of DMPC or DMPC/SM at 4:1 molar ratio further distinguished Aβ_1-40_’s secondary structures and aggregation kinetics as compared to respective lipid-nanodiscs (Fig.S4). Taken together, these findings suggest that Aβ_1-40_ aggregation in macro-nanodiscs is preferably modulated by the lipid bilayer surface as the large planar lipid bilayer enables lipid-peptide interactions with less influence of the polymer that forms the belt of the lipid-nanodisc. Thus, the studied PMA-macro-nanodiscs could be a useful tool to study membrane mediated amyloid aggregation.

**Table 1.** Thermodynamic parameters for Aβ_1-40_ binding to PMA nanodiscs.

| Nanodiscs | $\Delta H$ | $\Delta S$ | $T\Delta S$ |
| --- | --- | --- | --- |
| DMPC | -5.45 | 1.13 | 0.33 |
| DMPC:DMPG (4:1) | -6.64 | -2.02 | -0.60 |
| DMPC:DMPG (1:1) | -7.70 | -5.98 | -1.78 |

$\Delta H$ :kcal/mol; $\Delta S$ :cal/mol; $T\Delta S$ :kcal/mol; $K_d$ : $\mu M$

We next measured the binding affinity (K_d_) of Aβ_1-40_ with DMPC (100%) or DMPC mixed with a variable amount of an anionic lipid (DMPG) nanodiscs. Remarkably, the ITC measurements showed distinct thermograms for Aβ_1-40_ binding to different nanodiscs (Fig.2e and f). DMPC nanodiscs showed a micromolar (K_d_=56.3 μM) binding affinity to Aβ_1-40_ in solution (Fig.2e) which agrees well with a previous study^5^ that reported K_d_ <100 μM for zwitterionic liposomes. Nanodiscs containing 20 and 50% of DMPG showed different binding affinities of 37.6 μM and 45.7 μM, respectively (Figs. 2f and S5). The binding between Aβ_1-40_ and DMPC nanodiscs was favored by both negative enthalpy (-ΔH) and positive entropy (TΔS) contributions. On the other hand, Aβ_1-40_ binding to anionic nanodiscs was only favored by -∆H (Table 1). The increase in -∆H from -6.64 kcal mol^−1^ (4:1 DMPC:DMPG) to -7.70 kcal mol^−1^ (1:1 DMPC:DMPG) revealed a membrane dependent binding affinity for Aβ_1-40_ at a defined PMA concentration. In addition, the -∆S opposes Aβ_1-40_ binding to anionic nanodiscs which further supported its membrane selective binding. Overall, the ITC results suggest that Aβ_1-40_ interaction is membrane dependent although the cationic PMA triggers a uniform rapid β-sheet conformation in Aβ_1-40_ as revealed by CD.

To further examine the binding mechanism of Aβ_1-40_ with nanodiscs and its conformational states, we measured 2D ^15^N/^1^H SOFAST-HMQC NMR spectra of Aβ_1-40_ at 10 and 25 °C, and assigned the peaks based on a previous study.^17^ Poorly dispersed ^15^N/^1^H resonances obtained at 10 °C (Fig.3a, red) indicate a random-coil conformation which is in agreement with the CD results (Fig.2a). At room temperature (25 °C, incubated for 30 minutes in solution), we observed a slight change in the chemical shift and an increase in the peak intensity (Fig.3b, blue) for the amyloid core residues. The missing Serine and Glycine resonances in solution at 25 °C, were observed in the presence of nanodiscs (DMPC:DMPG=4:1) with 1:20 peptide:lipid molar ratio at 25 °C (Fig.3b, green). These observations suggest that Aβ_1-40_ bound to nanodiscs exhibits a short correlation time (τ_C_) and exists in a low-ordered aggregation state in comparison to its metastable amyloid fiber. The increase in peak intensity observed for residues S8, G9, K16, G25, S26 and V36 indicate the induction of a more flexible structure for the selected domains (Fig.S6). In contrast, the core domain comprising of residues 17-20 show a substantial decrease in peak intensity and suggests a membrane-associated and folded state.^7,18^ Taken together, the ThT aggregation kinetics that showed a T_lag_ over 20 hrs (Fig.1) for 4:1 DMPC:DMPG nanodiscs at 1:20 peptide:lipid molar ratio correlates with the dispersed ^15^N/^1^H NMR resonances, unlike that observed for matured fibers (Fig.S2c). This indicates the formation of low-ordered Aβ_1-40_ aggregates in presence of nanodiscs.

**Figure 3.**
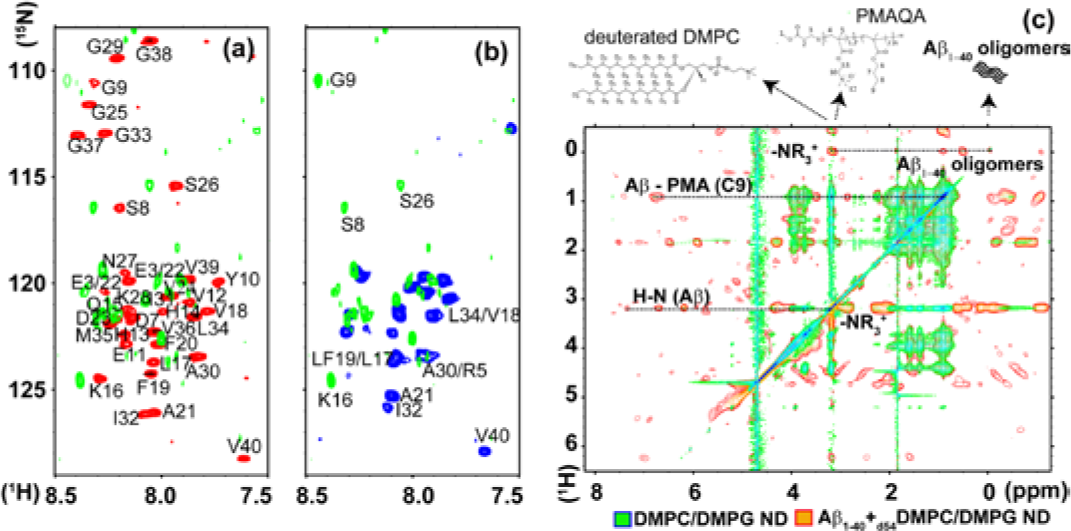
Structural and functional characterization of Aβ_1-40_ in lipid-nanodiscs. (a-b) ^15^N SOFAST-HMQC spectra of Aβ_1-40_ (70 μM) titrated with 1.4 mM DMPC:DMPG (4:1) nanodiscs in 10 mM sodium phosphate buffer solution (green). The red and blue spectra represent the peptide spectra acquired in absence of nanodiscs at 10 °C and 25 °C, respectively. (c) 2D NOSEY spectra of 70 μM monomeric Aβ_1-40_ in presence (red-yellow) or absence (green-blue) of 1.4 mM _d54_-DMPC/DMPG (4:1) nanodiescs at 25 °C. The dotted (horizontal) lines show the correlation peaks between the amide-protons of Aβ_1-40_ and the side chain resonances (c9 and-NR_3_^+^) of the PMA-belt in nanodiscs. chemical structures of _d54_-DMPC and PMA are shows at the top. NMR spectra were recorded on a 600 MHs Bruker NMR spectrometer using a cryoprobe.

2D ^1^H/^1^H NOESY spectrum showed two diagonal peaks close to 0 ppm indicating the presence of Aβ_1-40_ oligomers (Fig.3c).^19^ These oligomer peaks show cross-peaks with PMA’s hydrophobic chain (C9 atom) and the quaternary ammonium of polymer or lipids (Fig.3c). Both the amide (H-N) and side chain protons (γ) of Aβ_1-40_ show a substantial correlation with PMA functional groups (Fig.3c). The corresponding ^1^H NMR spectra of Aβ_1-40_ (Fig.S7b) showed a substantial dispersion in the amide region that are significantly different from Aβ_1-40_ monomers and fibers (Fig.S2b and c). The NOESY spectrum indicates a direct interaction between the quaternary ammonium group of PMA (in nanodiscs) and Aβ_1-40_.

**Figure 4.**
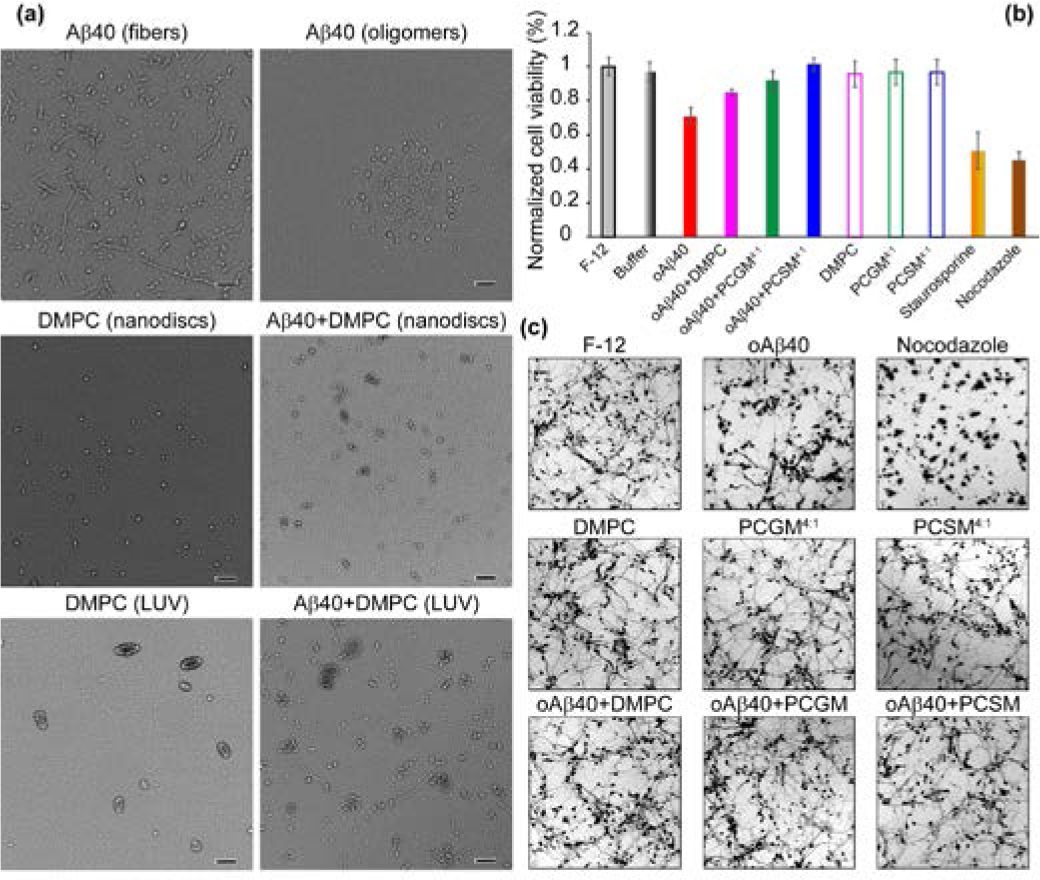
TEM and fluorescence images of nanodiscs and SH-SY5Y cells treated with Aβ_1-40_. (a) TEM images of Aβ_1-40_ fibers, oligomers and intermediates in presence of DMPC nanodiscs or LUVs measured after 48 hours. The scale bar is 200 nm. (b) MTT assays showing the formazan absorbance expressed as a measure of cell viability from SH-SY5Y cultured cells treated with 5 μM of Aβ_1-40_ oligomers (oAβ40) in presence and absence of nanodiscs. (c) Fluorescence images of neuronal damage caused by 5 μM of Aβ_1-40_ oligomers in presence and absence of nanodiscs. F-12 is the negative control; staurosporine and nocodazole were used as positive controls.

TEM images of Aβ_1-40_ fibers and oligomers, freshly prepared by dissolving monomers (see Methods in supporting information), showed morphologically distinct species after 48 hrs (Fig.4a). No Aβ_1-40_ fibers were observed in presence of DMPC nanodiscs (Fig.4a, middle). While nanodiscs fusion and aggregated spherical populations were seen in Aβ_1-40_ treated DMPC-nanodiscs (Fig.4a), a mixed fibril and spherical Aβ_1-40_ species were observed in presence of DMPC-LUVs (Fig.S3b). These observations demonstrate a successful trapping of lower-ordered aggregates/intermediates of Aβ_1-40_ species using nanodiscs (Fig.4a).

Protein-based nanodiscs have been shown to control the aggregation and toxicity of Aβ_1-40_ in-vitro and in-vivo^14,20–22^.

Thus, we studied the potential use of PMA-nanodiscs in modulating the toxicity of Aβ_1-40_ oligomers. The Aβ_1-40_ oligomers treated to human neuroblastoma (SH-SY5Y) cells were observed to be neurotoxic with a substantial reduction in cell-viability (Fig. 4b), and fluorescence imaging showed a severe damage to neurites and soma (Figs 4c and S8). Whereas, the Aβ_1-40_ oligomers incubated with PMA-nanodiscs exhibited relatively less neurotoxic and neuronal damage as shown in Fig.4b and c; the least toxicity observed for DMPC/SM nanodiscs is in agreement with the reported role of SM.^23^ Morphometric analysis of neurons treated with nanodisc containing Aβintermediates showed well-connected neurites that were severely damaged by oligomers and small molecules like staurosporine and nocodazole (Figs. 4c and S8). Aβ_1-40_ monomers or fibers interacting with nanodiscs showed a loss of ≈25% cell viability, whereas Aβ_1-40_ oligomers exhibited significant (≈60%) cytotoxicity (Fig.S9). Thus, we have demonstrated the PMA encased nanodiscs are potential molecules to protect neuron cells by reducing the toxicity of Aβ_1-40_ species.

Structural and mechanistic studies presented here revealed a symbiotic functional relationship for PMA and its associated membrane lipids. While PMA induces a rapid structural transition in Aβ_1-40_, a controllable intermediate of Aβ_1-40_ can be generated by varying the lipid composition and concentration in PMA-nanodiscs (Figs.1 and 2). In addition, as evidenced from NMR results that showed a dispersed ^1^H/^15^N resonance distribution and intermolecular ^1^H/^1^H correlation, polymer-nanodiscs can be used to track amyloid intermediates for real-time measurements (Fig.3). It is also remarkable to note that the “styrene-free” PMA-nanodiscs enable fluorescence and CD experiments for real-time characterization of amyloid aggregation and are useful for conformational analysis of amyloidogenic proteins embedded in a lipid bilayer.

While investigating the structure, dynamics and function of Aβ_1-40_ in membrane interface remains challenging due to the complexity, we have proposed the applicability of nanodiscs and macro-nanodiscs to decipher the molecular mechanism of seeding reaction at atomistic-scale on real-time. PMA encased lipid nanodiscs generates off-pathway lower-order aggregates of Aβ_1-40_ directed by PMA and modulated by lipid composition over time. Based on the NMR observation (Fig.3), we propose a distinct structural ensemble of Aβ_1-40_ intermediates. The toxicity of Aβ_1-40_ oligomers were substantially lowered by nanodiscs and are dependent on the lipid properties and yielded distinct morphological phenotypes as revealed from TEM and cell assay (Fig.4).^24^

In conclusion, we have successfully demonstrated a new approach to trap Aβ_1-40_ intermediates and characterize them at atomic-level using polymer nanodiscs for the first time. The significant binding efficacy of PMA-nanodiscs with Aβ_1-40_ species reported in this study could be valuable in the development of potential therapeutic strategies for Alzheimer’s disease. While the lack of understanding and high-resolution structures of membrane-bound Aβ_1-40_ intermediates have been a bottle-neck^25^, the structural and functional insights into Aβ_1-40_ intermediates reported in this study could also be valuable for the development of compounds to suppress neuronal cell death and potentially develop treatment for Alzheimer’s disease. The ability of polymer-based nanodiscs to trap amyloid intermediates and remodel them to pathologically distinct morphological states is unique and will have broad impacts in the structural characterization of a variety of membrane-assisted amyloid aggregation processes that are implicated in various neurodegenerative disorders. Further, this approach would also enable the application of a variety of physical techniques for high-throughput characterization of protein misfolding, nanodisc encapsulated AD drug delivery and screening of small molecule compounds for the development of therapeutics. While the PMA-nanodiscs render a curvature-free lipid bilayer to probe the role of the lipid membrane on amyloid aggregation, further investigation to fully understand the mechanism of the polymer-belt induced formation of β-sheet structure of Aβ and the use of PMA-nanodiscs on other amyloid proteins would be useful. Further development of different types of polymer-belts like peptide and apolipoproteins-encased nanodiscs^14,22^ for the stabilization of amyloid oligomers would be also beneficial for high resolution structural studies.

## Acknowledgements

This study was supported by NIH (AG048934 to A.R.). We thank Professor Bernd Reif for providing us the recombinant expression system and protocol for the production of amyloid-beta-1-40 peptide.

